# Estrogen suppresses DMRT1 expression during ovarian development in the chicken

**DOI:** 10.1101/2022.02.22.481469

**Authors:** Debiao Zhao, Long Liu, Sunil Nandi, Jason Ioannidis, Xiurong Yang, Daoqing Gong, Mike J. McGrew, Michael Clinton

## Abstract

Primary sex determination is the choice between two different developmental fates, a testis or an ovary. This selection is thought to require the action of a master regulator gene that triggers expression of a gene cascade in the bipotential gonad precursor in one sex. The selection of a particular developmental pathway is also thought to suppress the alternative developmental pathway.

In birds, where the male is the homogametic sex (ZZ) and females the heterogametic sex (ZW), the Z-linked transcription factor DMRT1 is considered the master regulator and has been shown to be essential for testis development, and to also inhibit the ovarian pathway. Here, we characterise in detail, DMRT1 transcription and protein levels during gonadal development in the chick. Our analysis suggests that DMRT1 protein levels are equivalent in male and female gonads during the bipotential phase of development, and that DMRT1 protein levels are reduced in the developing ovary during the differentiation phase. The reduction in DMRT1 protein levels in the somatic cells of the female medulla, coincides with the induction of aromatase expression and the initiation of estrogen synthesis. Analysis of sex-reversed gonads and mixed-sex chimeric gonads, suggests that the reduction in DMRT1 protein is due to inhibition of DMRT1 expression by estrogen.

Our data suggests that estrogen signalling is involved in primary sex determination by regulation of DMRT1 protein expression.

## Introduction

While the genetic factors that regulate gonadal differentiation in vertebrates are generally well conserved, the factors that initiate primary sex determination (development of the gonads) differs between different species. In the most-studied of vertebrates, the mammal, expression of the Y chromosome encoded *SRY* gene in the male gonadal primordia (genital ridges), induces testicular differentiation, while in females which lack the *SRY* gene, a different set of genetic factors induces ovarian differentiation (reviewed in 1). Examples of male differentiation factors used by other vertebrate species include DMRT1 (Doublesex and Mab-3 Related Transcription factor 1) and AMH (anti-Mullerian hormone). These factors are utilised in different ways in different species (2): for example, fishes employ a variety of sex-determining genes, including *dmrt1*, *dmrt1y*, *sdy*, *amhy* and *amhr2*. Some amphibians and reptiles use *Dmrt1* homologs and paralogs, such as *dmw*, and sometimes under the control of external stimuli (3–6). Although primary sex determination in mice and humans is not driven by DMRT1, it does play a role in maintaining male somatic cell sex identity in adult testes (7,8).

In birds, the male is the homogametic sex (ZZ) and the female is the heterogametic sex (ZW), and previous research has suggested that primary sex-determination in birds is likely to depend on a gene dosage mechanism based on a Z chromosome gene(s) (9), with the most likely candidate, the Z chromosome *DMRT1gene* (10–13). Recently we have conclusively shown that *DMRT1* is the key factor in primary sex determination in chickens (14).

Using a gene editing approach, we demonstrated that the presence of two functional copies of the *DMRT1* gene promotes testis development in male chick embryos. During development, male chick embryos develop two paired testis while female embryos develop only a single ovary on the left side. We showed that the loss of a single copy of *DMRT1* led to the male embryo developing gonadal architecture typical of a female, i.e. the development of a single gonad – an ovary – on the left side. Despite this advance, the mechanism whereby *DMRT1* regulates cell fate in embryonic chick gonads is unclear. It is widely assumed that this dosage-based mechanism is dependent on higher levels of active transcription factor in males, and numerous studies have supported this theory by showing that, although *DMRT1* is expressed in the developing gonads of both males and females, total transcript levels are higher in males than in females (12,15). A key study also demonstrated that suppressing levels of *DMRT1* transcript in male gonads results in the ‘feminisation’ of certain medullary cells (13). The assumption is that higher levels of transcript correspond to higher levels of active DMRT1 protein in cells in male gonads than in the equivalent cells in female gonads, but this has yet to be effectively demonstrated. An alternative possibility is based on the fact that the *DMRT1* locus generates a number of different transcripts (16, Suppl Fig 1) and so it is possible that different transcripts are expressed in male and female gonads. It is also not clear that the medullary cells that express *DMRT1* in male and female gonads are equivalent. In addition to certain somatic cell types, *DMRT1* is also expressed by germ cells, raising questions as to whether sex differences in transcript levels identified in ‘whole gonad’ analyses, accurately reflect differences between the somatic cells of male and female medullas.

For these reasons we have elected to review expression of DMRT1 in the developing male and female chick embryo gonads. This analysis has shown that DMRT1 protein levels do not directly reflect *DMRT1* transcript levels in male and female gonads. Our results suggest that DMRT1 protein levels are similar in male and female gonads in the initial stages of gonadogenesis, and that the sex differences in DMRT1 protein levels seen at later stages of gonadal development, are due to the suppression of DMRT1 expression in females rather than elevated levels of DMRT1 expression in males. Our analysis also suggests that this suppression of DMRT1 in the developing ovary, is co-incident with, and may be a result of, rising levels of estrogen.

## Results

First, we investigated whether the same *DMRT1* transcripts were expressed in male and female gonads. We isolated poly-A RNA from pools of male gonads and pools of female gonads collected at different stages of embryonic development between E4.5 and E8.5. This material was used for Northern analysis and hybridised against a cDNA probe containing the region corresponding to the DM domain of *DMRT1* (Figure 1a). This analysis revealed a single transcript of 1.3 kb in length present in both male and female samples. This transcript corresponded to transcript isoform b and was expressed in all male and female samples between E4.5 and E8.5 (the period of sex determination). It is clear that this *DMRT1* transcript is present at higher levels in male than in female samples at all stages studied, and also that expression levels in male gonads and in female gonads, vary little over this key developmental period.

**Figure 1.**
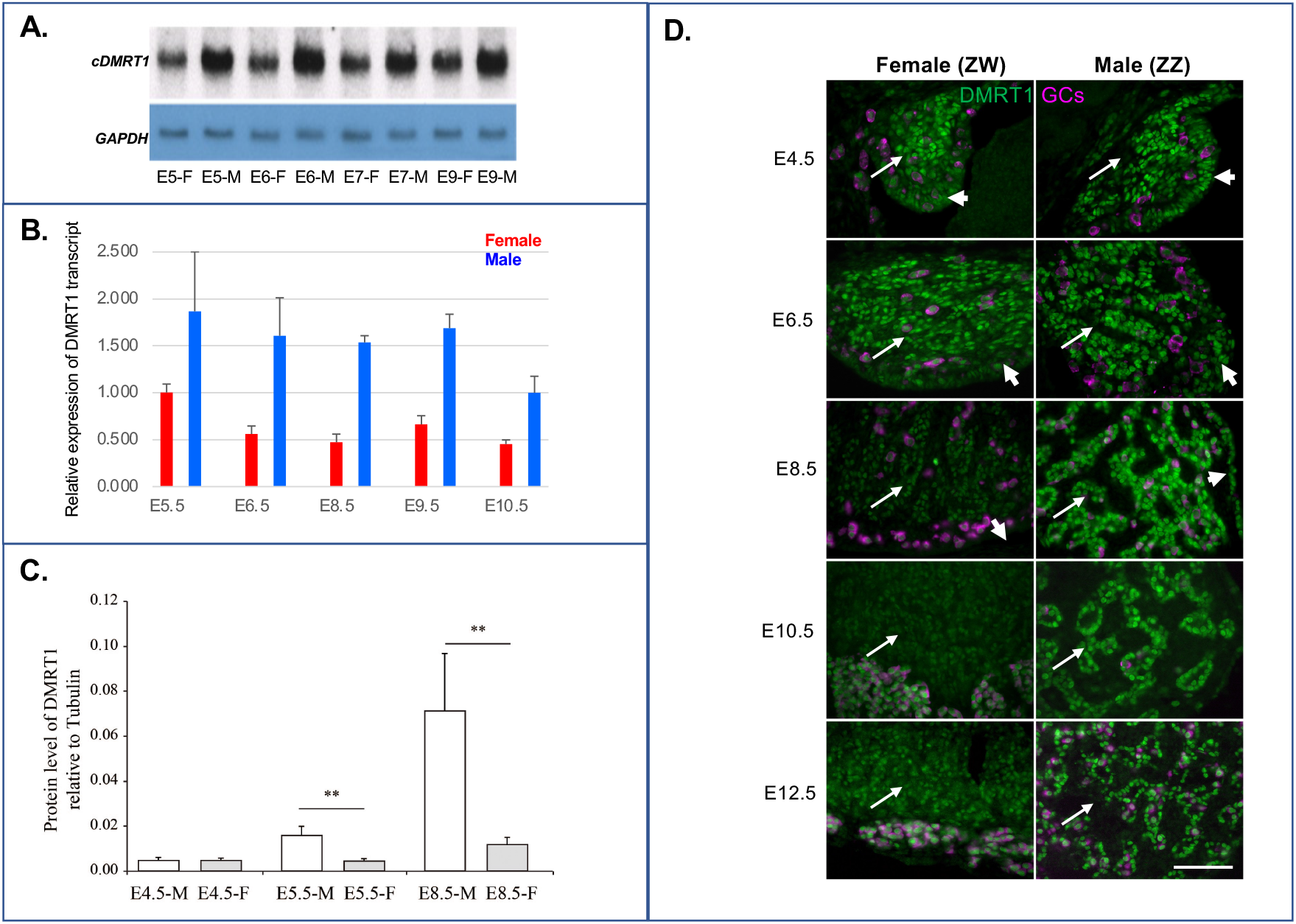
Expression of DMRT1 transcript and protein in the developing gonads of male and female chicken embryos. A. PolyA RNA Northern analysis of DMRT1 expression in female and male gonads from embryonic day 5 to embryonic day 9 (E5 – E9). For each sample, PolyA RNA was isolated from 60mg of gonad/mesonephros total RNA by oligo dT selection. Northern membrane was hybridised sequentially with radiolabelled cDNA probes for DMRT1 and GAPDH. DMRT1 transcript length was 1.3kb. (F=female; M=male). B. qRT-PCR analysis of DMRT1 transcript levels in female (red) and male (blue) gonads from E5.5 to E10.5. DMRT1 levels were corrected for loading variation by comparison to GAPDH levels and are show relative to DMRT1 levels in E5.5 female gonads. Gonad pairs were pooled and a minimum of 5 individual pools of each sample type were analysed. C. Example of Western analysis of DMRT1 protein levels in male and female gonads between E4.5 and E8.5. DMRT1 levels were corrected for loading variation by comparison to b-Tubulin levels. (** indicates significant difference (P<0.01); gonads were pooled for analysis and 2 individual pools were analysed for each sample type). D. Immuno-Histochemistry (IHC) analysis of DMRT1 protein expression in different cell types during gonadal development. Histological sections of female and male gonads between E4.5 and E12.5 of development were incubated with antibodies against DMRT1 and the germ cell (GC) protein VASA. Arrows indicate gonadal medulla and arrowheads indicate coelomic epithelium. (scale bar indicates 50mM).

Having established that only a single transcript isoform is expressed in the developing gonads, we carried out a qPCR analysis of *DMRT1* expression to more accurately quantitate relative male:female differences. We measured *DMRT1* levels in male and female gonads between E5.5 and E10.5 of development. Any variation in input cDNA levels was corrected by comparison to *GAPDH* transcript levels and *DMRT1* levels are displayed relative to *DMRT1* expression in E5.5 female gonads (Figure 1b). It is clear that, in agreement with previous reports, that levels of the DMRT1 transcript are at least 2-fold higher in whole male gonads than in whole female gonads throughout this period. It is also clear that levels of *DMRT1* transcript in male gonads do not vary significantly between E5.5 and E9.5, while levels in female gonads decrease from E6.5.

To assess whether transcript levels reflected protein levels, we examined DMRT1 protein levels in male and female E4.5, E5.5 and E8.5 gonads by Western analysis (Figure 1c). Although *DMRT1* transcript levels were consistently 2-fold higher in whole male gonads than in whole female gonads, Western analysis revealed a different profile for DMRT1 protein levels. At E4.5, DMRT1 protein levels were similar in male and female whole gonads; at E5.5, DMRT1 protein levels were approximately 2-fold higher in male gonads than in female gonads; and at E8.5, DMRT1 protein levels were approximately 5-fold higher in male than in female gonads. As with *DMRT1* transcript analyses, the Western analysis reflects DMRT1 protein levels not only in the somatic cell of the medulla, but also in the surrounding epithelium/cortex and in the germ cells.

To examine the cellular location of DMRT1 protein we performed immunohistochemistry on histological sections from male and female gonads at five stages of development between E4.5 and E12.5 (Figure 1d). Sections are from the left gonad in all instances and show the gonadal medulla, the overlying epithelium and the germ cells. At E4.5, nuclear DMRT1 is evident in all three tissue compartments, with highest levels seen in the somatic cells of the medulla. At this stage of development, the signal intensity and tissue distribution is similar in both male and female gonads. At E6.5 the tissue pattern of expression is similar to that of E4.5, although the immunofluorescence signal intensity appears to be slightly higher in the somatic cells of the male medulla than in the corresponding cells of the female medulla. By E8.5, there is a marked difference in DMRT1 protein levels and tissue patterns in male and female gonads. In male gonads, DMRT1 protein is restricted to the cells of the developing sex cords (somatic and germ cells) and to the surrounding epithelium. Medullary cells outwith the sex cords do not express DMRT1. In the female gonad at E8.5, DMRT1 protein is present in cells throughout the medulla and in the germ cells. At E8.5, DMRT1 protein levels in the somatic cells of the sex cords in the male gonad are significantly higher than in DMRT1-expressing cells in the female gonad. The expression patterns seen at E10.5 and E12.5 are similar to that at E8.5: high levels of DMRT1 protein in the sex cords of the male medulla and lower levels in cells of the female medulla. In the female gonad, DMRT1 protein levels are highest in the germ cells that have relocated to the developing ovarian cortex. It is interesting to note that the difference in DMRT1 protein levels between male and female gonads appears to be due to a decrease in DMRT1 protein levels in the female gonad from E6.5 onwards, rather than an increase in DMRT1 protein levels in the male gonad.

Our analyses suggesting that DMRT1 protein levels are similar in male and female gonads during the initial stages of gonadogenesis, conflicts somewhat with accepted dogma. Previous reports, and assumptions based on gene transcript levels (including our own data), suggested that DMRT1 protein levels would be significantly higher in male gonads than in female gonads at all stages of development, whereas the current study suggests that this is not the case. However, when comparing individual IHC sections, although all samples are processed concurrently, it is still possible that observed differences in signal intensity between male and female sections are artifactual and due to variation in tissue processing and/or in staining procedures. In order to exclude this possibility, we generated embryos with mixed-sex chimeric gonads; i.e. gonads composed of a combination of male and female elements. In this model, undifferentiated mesoderm is transplanted from donor to host embryo at a very early stage of development and undergoes proliferation, migration and differentiation alongside host tissue, in the development of the gonads (17). The resulting gonads are comprised of a combination of genetically male and genetically female somatic tissues (germ cells are derived from host embryo) (Figure 2A). Of course, when histological sections of these gonads are used for IHC, the male and female compartments will have been subjected to identical processing and staining procedures. We generated embryos with mixed-sex chimeric gonads and compared DMRT1 protein levels in male and female portions at E5.5 and at E8.5. In our approach, the donor embryos ubiquituously express GFP, enabling us to distinguish between donor and host tissues. In the examples shown, tissue was transferred from female GFP embryos into wild-type male embryos at E2.5, and eggs sealed and re-incubated until either E5.5 or E8.5 (Figure 2). The donor portions of the resulting gonads are easily identified by GFP expression. It is clear from this analysis that the DMRT1-expressing cells in the male and female portions of this E5.5 chimeric gonad, display similar signal intensities, suggesting that somatic cells of the male and female portions of the medulla, express similar levels of DMRT1 protein at E5.5 (Figure 2 b.ll). In contrast, a similar analysis of male:female chimeric gonads at E8.5, suggests that DMRT1 protein levels are higher in the somatic cells of the male portion than in the somatic cells of the female portion (Figure 2 b.lV). This analysis of mixed-sex chimeric gonads appears to confirm our original finding that, during the early stages of gonadal development in the chick, DMRT1 protein levels are similar in the medullary somatic cells of male and female embryos.

**Figure 2.**
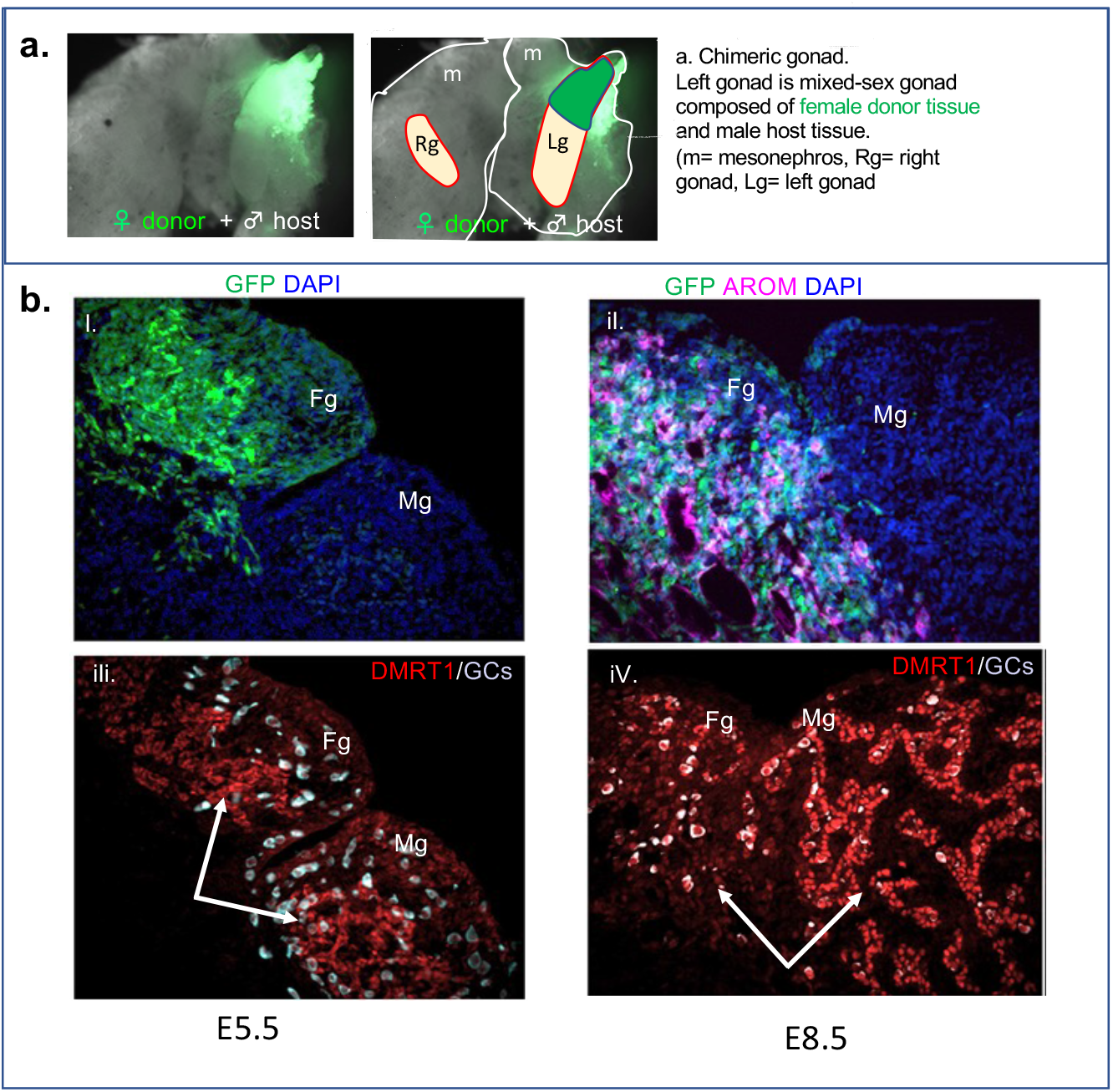
Expression of DMRT1 protein in female and male regions of mixed-sex chimeric gonads. a. Gross morphology of mesonephroi and gonads from mixed-sex chimera. Panel on right shows schematic illustration of host and donor regions. b. IHC analysis of protein expression in histological sections of mixed-sex chimeric gonad removed at E5.5 and E8.5 of development. Arrows indicate DMRT1-expressing cells in medullary regions of female and male portions of mixed-sex gonad. Green Fluorescent Protein (GFP) indicates female donor tissue (AROM=aromatase; GCs= germ cells; Fg= female donor derived region; Mg= male host region; DAPI staining shows individual cell nuclei).

We have recently confirmed that *DMRT1* gene dosage is a key factor in primary sex-determination in birds (14). This study showed that the loss of a functional copy of *DMRT1* in male embryos resulted in ovarian rather than testicular development, and also that this transformation was accompanied by the induction of FOXL2 expression in cells of the male gonadal medulla. To explore the relationship between DMRT1 and FOXL2, we directly compared expression of both proteins in the developing ovary (Figure 3). In the initial stages of ovarian development, DMRT1 protein is present at high levels in the majority of the cells of the medulla, while the expression of FOXl2 is restricted to a limited number of cells in the ventral portion of the medulla, underlying the coelomic epithelium. As development proceeds, the proportion of medullary cells expressing FOXL2 protein increases, while the level of DMRT1 protein in these cells decreases. Expression of FOXL2 appears to spread from the ventral region of the medulla to the dorsal region. By E8.5, the gonad has assumed an ovarian appearance, with a thickened cortex surrounding the medulla core. By E9.5, DMRT1 is uniformly expressed in cells throughout the medulla but at levels lower than those seen in the early stages of gonadogenesis. By this stage, all DMRT1-expressing somatic cells of the medulla also express FOXl2. The increase in FOXL2 protein expression in the medullary somatic cells during ovarian development is clearly concurrent with a reduction in DMRT1 protein levels. This process initiates in the ventral region of the gonad immediately adjacent to the developing cortex and extends across the entire medulla between embryonic stage 5.5 and E8.5.

**Figure 3.**
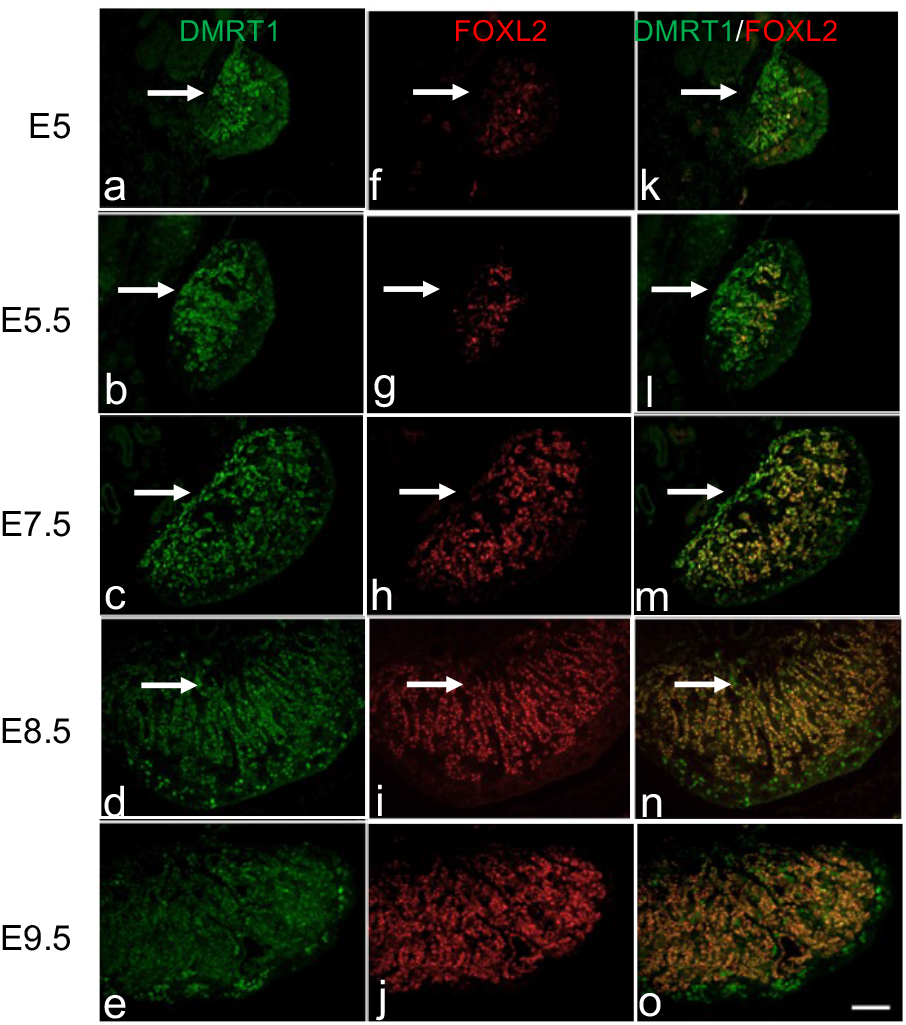
Expression of DMRT1 and FOXL2 proteins in the developing chick ovary. IHC analysis of DMRT1 and FOXL2 expression in the left female gonad between E5 and E9.5 of development. Images illustrating DMRT1 expression are shown in panels a-e and images illustrating FOXL2 expression are shown in panels f-j. Merged DMRT1/FOXL2 images are shown in panels k-o. Arrows indicate dorsal region of gonadal medulla. Scale bar indicates 50 mM.

Our findings suggest that, during ovarian development, the induction of FOXL2 is accompanied by a suppression of DMRT1 expression. FOXL2 has been shown to induce expression of the enzyme aromatase (18–20) which converts androgens to estrogens. Ovarian cortex formation is dependent on the synthesis of estrogen by aromatase-expressing cells of the medulla, and it is possible that estrogen is responsible for suppressing DMRT1 expression. To explore this possibility, we examined expression in experimentally induced sex-reversed gonads. It is well established that treatment with the chemical fadrozole which inhibits aromatase activity, induces ovary-to-testis sex reversal in female chicken embryos (21–23). By moderating the dosage of fadrozole we generated female embryos with gonads in transition from ovary to testis (ovotestis). Figure 4 shows IHC analysis of sections from the left gonads at E9.5 from male, female and partially sex-reversed female embryos. This figure clearly shows the presence of sex cords in the developing testis (K & l), an obvious cortex in the developing ovary (a - c) and a marked difference in DMRT1 protein levels in the male and female medullas (k & b). There is no cortex present in the fadrazole-treated female gonad (d – i), indicating a lack of estrogen. The ovo-testis nature of the gonad from the fadrazole-treated female embryo, is illustrated by expression of the male marker SOX9 in the dorsal region of the medulla (e) and expression of aromatase in the ventral region of the medulla (d & g). The SOX9-expressing dorsal medulla shows clear signs of sex cord development typical of a testis, while the ventral aromatase-expressing region is largely unstructured. It is clear that the dorsal region of the medulla expresses higher levels of DMRT1 protein than the ventral region (h). This analysis suggests that the loss of estrogen in the female gonad coincides with an increase in DMRT1 expression (from the single Z-chromosome), and with the loss of aromatase.

**Figure 4.**
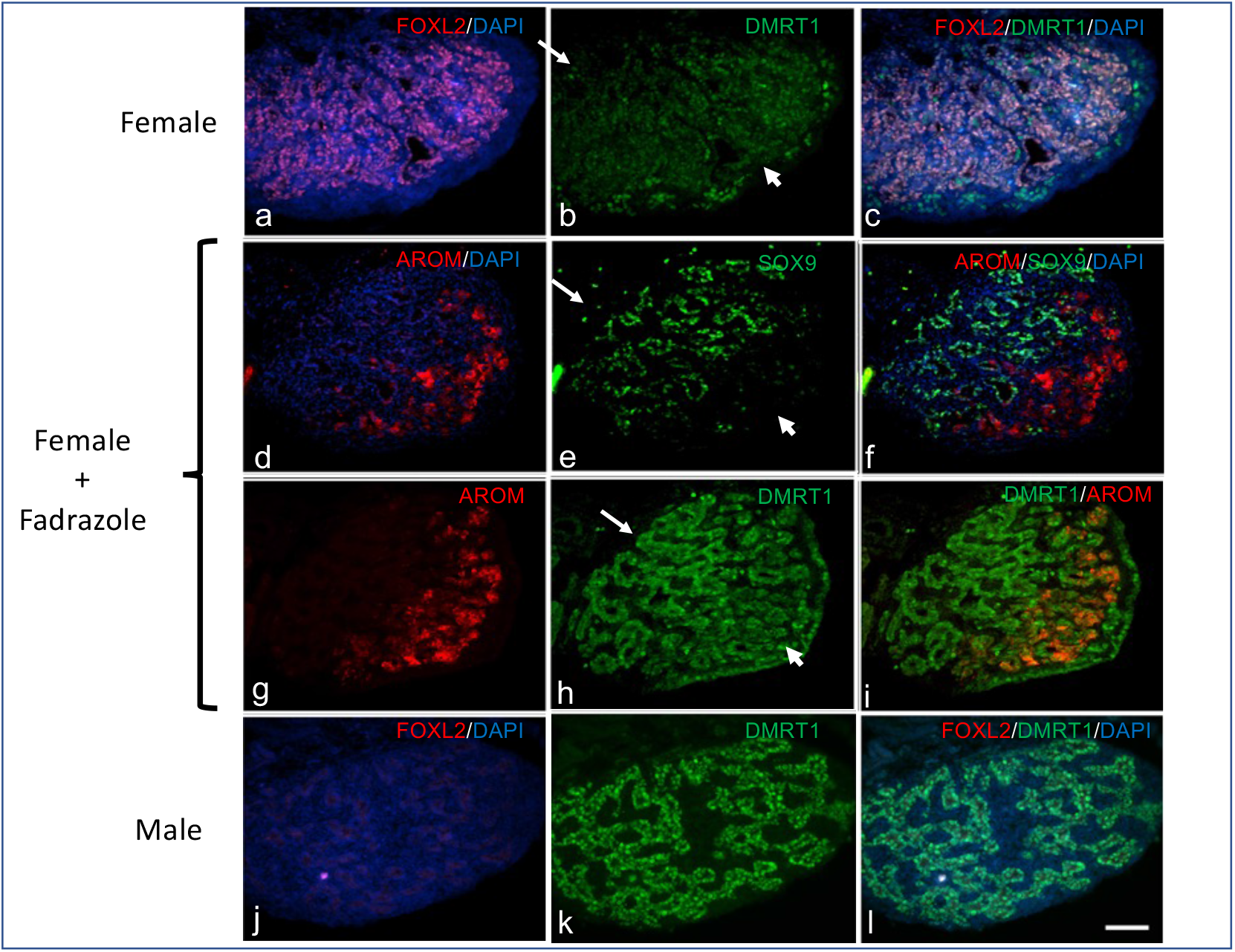
Expression of female and male marker genes in left gonads of untreated and Fadrazole-treated embryos. IHC analysis of DMRT1, SOX9, FOXL2 and aromatase (AROM) protein expression in histological sections from female (panels a-c) and male (panels j-l) gonads, and in Fadrazole-induced ovo-testis (panels d-i) at E9.5. Sections shown in panels d-f and in panels g-I, represent adjacent tissue sections. Scale bar indicates 50 mM. DAPI indicates nuclear staining. Arrows indicate dorsal region of gonadal medulla; arrowheads indicate ventral/epithelium surface of gonad.

To further explore this relationship we utilised an alternative means of inhibiting estrogen signalling. Fulvestrant is an anti-estrogen that blocks estrogen signalling by binding to, and inducing degradation of the estrogen receptor (ER) (24). We treated chicken embryos at E2.5 of development with either fulvestrant alone, or fulvestrant in combination with the aromatase-inhibitor, fadrozole (Figure 5). At the gross morphological level, fulvestrant treatment alone appeared to cause complete ovary to testis gonadal sex reversal in female embryos. IHC analyses of histological sections revealed that the medulla of the left gonad was composed predominately of SOX9-expressing sex cords (g-j), and that there was no obvious cortical tissue present. A limited region close to the ventral surface retained aromatase expression. The left gonad of female embryos treated with a combination of fulvestrant and fadrozole displayed a similar profile. DMRT1 protein levels were compared in the gonads of untreated male and female embryos and the gonads of embryos treated with either fluvestrant alone or fluvestrant in combination with fadrozole. It is clear that the signal intensity and pattern of DMRT1 expression in the gonads from both types of treated embryos are similar to those in the wild-type male gonads, and distinct from those in wild-type female gonads. This data suggests that blocking estrogen signalling can alleviate the suppression of DMRT1 protein expression seen during ovary development.

**Figure 5.**
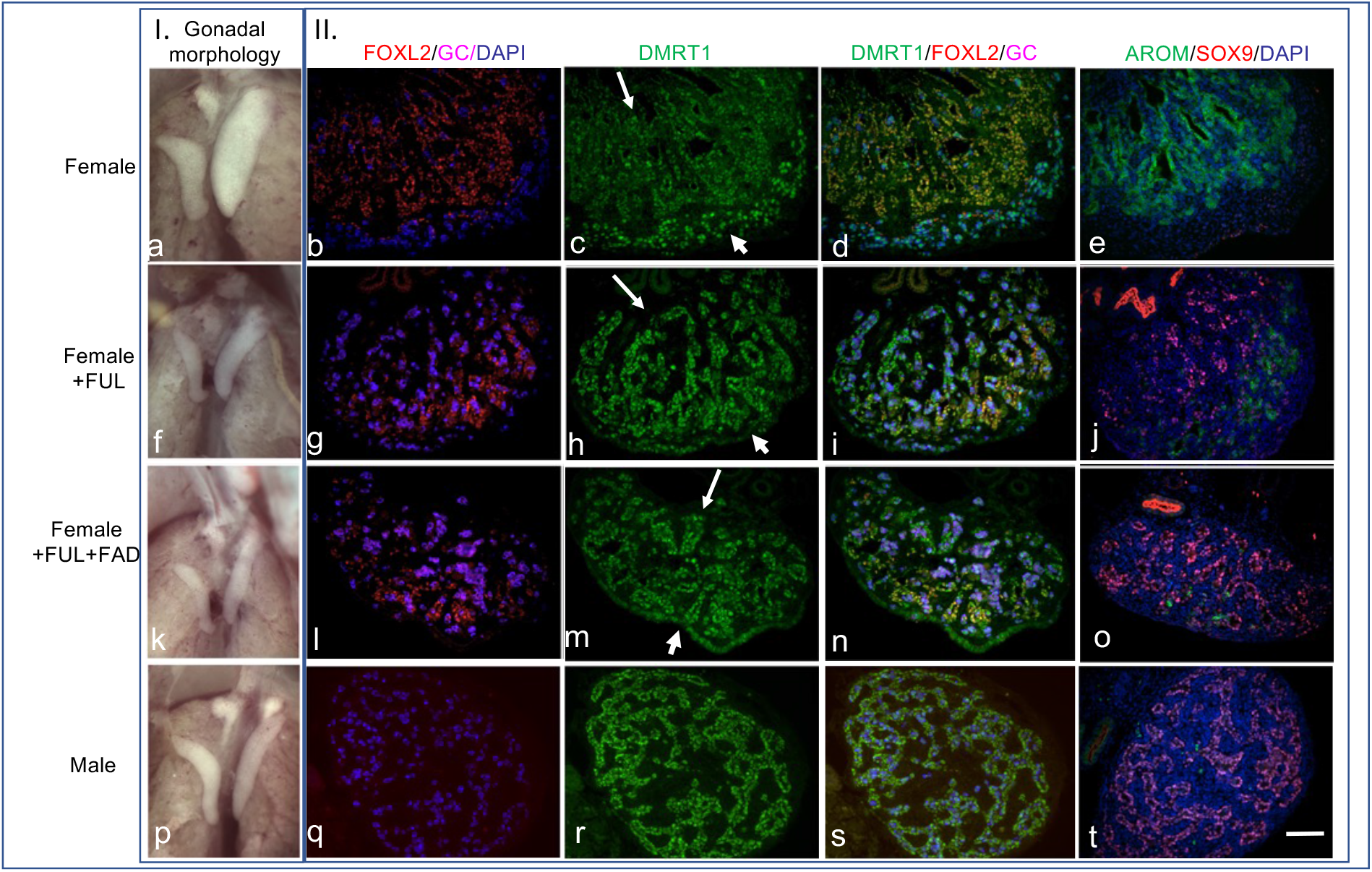
Gonadal morphology and marker gene expression in left gonad of untreated and sex-reversed embryos. I. Light microscopy images of gonads and mesonephroi from female (a), male (p) and sex-reversed embryos (f, k) at E9.5 of development. II. IHC analysis of histological sections from left gonads of female (b-e), male (q-t) and sex-reversed embryos (g-o) at E9.5 of development. Scale bar indicates 50 mM. DAPI indicates nuclear staining. Arrows indicate dorsal region of gonadal medulla; arrowheads indicate ventral/epithelium surface of gonad. (FUL=Fluvestrant; FAD= Fadrazole).

Finally, we compared Estrogen Receptor-alpha (ER) expression in the gonads of male and female embryos (Figure 6). ER is detectable in both the somatic cells of the medulla and the coelomic epithelium from E7 of development (25,26). When bound by estrogen, the ER translocates to the nucleus (27) and can be visualised by IHC. To exclude the possibility of IHC-generated artifacts and to directly compare ER activation in male and female gonads, we analysed expression in mixed-sex gonads at E8.5 of development. These chimeric gonads are comprised of a region of male tissue and a region of female tissue, and estrogen production is dependent on aromatase activity in the medulla portion of the female component (see Figure 2). Our analysis shows that ER is present in the nuclei of the female portion of the chimeric medulla (low DMRT1/ aromatase +ve), but not in the nuclei of the male portion of the chimeric medulla (DMRT1 high/ aromatase -ve) (Figure 6d). However, our analysis also shows that ER is activated (nuclear) in the epithelium surrounding the medulla in both male and female portions of the chimeric gonad (Figure 6b & d). It is noteworthy that the epithelium expresses low levels of DMRT1 throughout development. As the IHC antibody can not detect cytoplasmic ER in the female portion of the chimeric medulla, it is not clear whether ER is not activated or simply not present in the male portion of the chimeric medulla.

**Figure 6.**
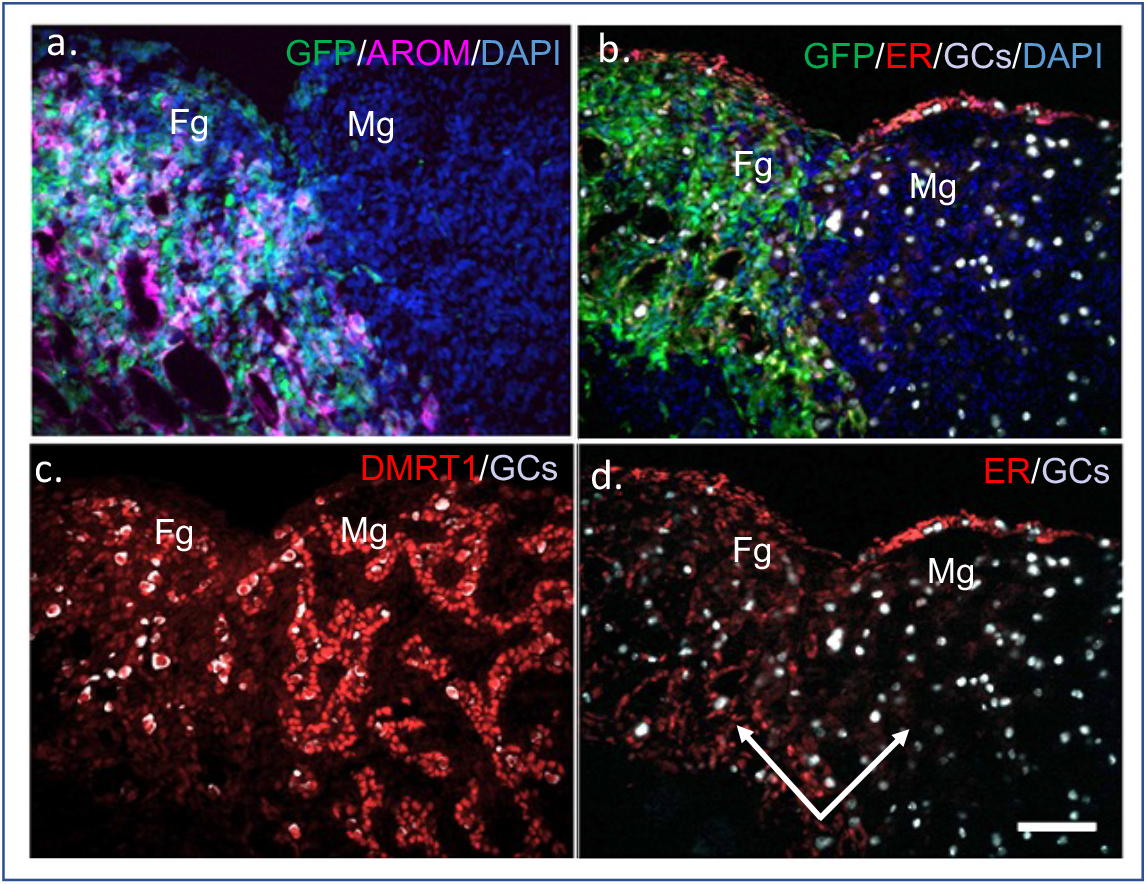
IHC analysis of aromatase (AROM), ER and DMRT1 protein expression in histological section of mixed-sex chimeric gonad at E8.5 of development. Panels a and b indicate female donor portion of chimeric gonad by expression of aromatase and Green Fluorescent Protein (GFP). Panel c replicates Figure 2 and illustrates that there are higher levels of DMRT1 protein present in the medullary cells of the male portion of the tissue than in the female portion. Panel d shows that ER is expressed and activated in both the medulla and the epithelium of the female portion, but only in the epithelium of the male portion of the chimeric gonad. Arrows indicate medullary cells in female portion and in male portion. GFP indicates presence of female donor tissue – host tissue is male. Fg= female portion of chimeric gonad; Mg= male portion of chimeric gonad, GC= germ cells. Scale bar indicates 50 mM.

## Discussion

The gonad primordium is considered a unique tissue on the basis that it has the capacity to differentiate into either of two distinct organs. Primary sex determination initiates a process whereby the undifferentiated gonad develops as either a testis or an ovary. The selection of a particular developmental pathway is also thought to suppress the alternative developmental pathway. Birds differ from mammals in that the female is the heterogametuc sex (ZW) and the male is the homogametic sex (ZZ). Attempts to identify a W-linked ovary-determining gene have been unsuccessful, and evidence has accumulated that the Z-linked DMRT1 gene is a testis-determining gene in birds. Over-expression of DMRT1 in females leads to masculinisation of the gonads while suppressing expression of DMRT1 in males leads to feminisation of the gonads (11,13). This sex-determining mechanism is thought to be based on gene dosage: expression of DMRT1 is not ‘dosage compensated’ and consequently, expression levels in males are higher than levels in females. A recent gene-editing approach has confirmed that two functional copies of the Z-chromosome *DMRT1* gene are required for testis development in the chicken (14). Although *DMRT1* is clearly a key factor in primary sex determination in chickens, the exact mechanism underlying this process is unclear, and may not be as straightforward as relative levels of this transcription factor. For this reason, we carried out a detailed examination of *DMRT1* transcription and protein expression during gonadal development in the chicken.

Here we examined DMRT1 transcription by Northern and q-PCR approaches. These analyses excluded the possibility that the sex determination mechanism depended on expression of different alternatively-spliced DMRT1 transcripts in males and females, and also confirmed that DMRT1 transcription was consistently higher in male than in female gonads. In the initial stages of gonadal development, DMRT1 expression levels in male gonads are approximately two-fold higher than expression levels in female gonads. During testes differentiation, initial DMRT1 levels were maintained, whereas during ovarian differentiation, DMRT1 expression levels decreased significantly. This supports the theory that the sex-determining effect of *DMRT1* is due to differential expression in male and female gonads, which leads to higher levels of this transcription factor in male gonads than in female gonads.

Western analysis of DMRT1 protein in the developing gonads, confirmed that higher levels of *DMRT1* transcription results in higher levels of DMRT1 protein, with the exception of E4.5 gonads where DMRT1 protein levels are similar in male and female gonads. However, it is difficult to make definitive conclusions on relative male/females levels by this approach, as in the later stages of development the male and female gonads are morphologically distinct. To enable comparison of DMRT1 protein levels in individual cell types, we carried out an IHC analysis of DMRT1 expression in male and female gonads between E4.5 and E12.5. At the early stages (E4.5 – E6.5), DMRT1 protein is present in all cells of both the male and female gonads, and at higher levels in the medulla than in the epithelium. In early gonadal development, DMRT1 protein levels appear to be similar in the somatic cells of both male and female gonads. However, by E8.5 there is an obvious difference in DMRT1 protein levels in male and female gonads: in the somatic cells of the male medulla, DMRT1 protein is expressed at higher levels than in cells of the female medulla. These differences in male and female DMRT1 protein expression are maintained at E10.5 and E12,5. Between E8.5 and E12.5, male/female difference in protein levels seem to correlate with the male/female difference in *DMRT1* transcription levels reported here and elsewhere. However, our results differ from those expected in that the male/female difference in protein levels seems to be due to a decrease in protein levels in female gonads rather than an increase in male gonads i.e. signal intensity is similar in male and female gonads in the early stages and while similar levels are maintained in the male gonads at later stages, levels decrease in female gonads. Analyses of gonads from mixed-sex chimeras supported this conclusion. In general, it is assumed that a double gene dose of *DMRT1* leads to higher levels of *DMRT1* transcription and higher levels of DMRT1 protein in developing male gonads than in developing female gonads, and that this induces testicular development in male embryos. Our data would appear to conflict with this assumption: it suggests that DMRT1 protein levels are similar in male and female medullas in the early stages of gonadal development, and that the suppression of DMRT1 protein leads to ovarian development.

As we had previously shown that loss of a single functional copy of *DMRT1* leads to the induction of FOXL2 in the developing gonads of male embryos, we compared the expression of DMRT1 and FOXL2 proteins in the developing left ovary of female embryos. Our analysis showed that in the early stages of ovarian development, when the somatic cells of the medulla display ‘male-like’ levels of DMRT1 protein, expression of FOXL2 protein is restricted to a narrow area immediately below the epithelium on the ventral side of the gonad. As ovarian development progresses, DMRT1 protein levels decrease in a wave from the ventral aspect towards the dorsal aspect of the medulla, and this is complemented by a ventral-to-dorsal wave of increasing FOXL2 protein. FOXL2 has been shown to induce aromatase expression which in turn catalyses the conversion of androgens to estrogens. The increase in FOXL2 expression that we find, coincides with an increase in estrogen synthesis (28), raising the possibility that estrogen suppress DMRT1 expression and enables ovarian development. To investigate this possibility, we manipulated estrogen levels in embryos and examined the effect on DMRT1 expression in the gonads.

Blocking aromatase activity or inhibiting estrogen signalling in the developing ovary led to gonadal sex-reversal with a decrease in aromatase expression, an increase in DMRT1 protein levels, and the formation of SOX9 +ve sex cords. DMRT1 protein levels appeared to be higher in the dorsal aspect than in the ventral aspect of the gonad: in partially sex-reversed gonads, the ventral medulla contained FOXL2 and aromatase +ve cells expressing low levels of DMRT1protein, while the dorsal medulla contained SOX9 +ve cells expressing high levels of DMRT1 protein. These findings are consistent with reports on other nonmammalian vertebrates, where exposure to estrogen results in male-to-female sex-reversal and inhibiting estriogen leads to female-to-male sex-reversal (29–35). In many instances, manipulating estrogen levels affects expression of DMRT1.

We also examined activation of estrogen signalling in the E8.5 mixed-sex chimeric gonads. In these chimeric gonads, aromatase is expressed in the somatic medullary cells of the female portion. We found that ER is activated (nuclear) in regions displaying low levels of DMRT1 protein (female medulla and both male and female epithelium) and not activated in regions with higher levels of DMRT1 protein (male medulla). It is well established that the epithelium of the male gonad can be induced to form an ovarian cortex in the presence of estrogen, and here we find that the ER is activated in that tissue.

This study provides a detailed characterisation of DMRT1 expression during gonadal development in the chick, and compared DMRT1 expression with FOXL2 expression and estrogen signalling. Our findings support the hypothesis that DMRT1 is the key factor in avian sex determination; promoting Sertoli cell differentiation and testis development and inhibiting FOXL2 expression and ovarian development in the male. Contrary to expectations, our findings suggest that during the initial stages of gonadogenesis, DMRT1 protein levels in female gonads are equivalent to levels in male gonads. We suggest that during ovarian development, in addition to inducing cortex expansion, estrogen signalling suppresses DMRT1 expression and prevents Sertoli cell differentiation and sex cord formation in the medulla. This is supported by the findings of an analysis of male birds with a single functional copy of DMRT1 (14). These birds develop ovaries in place of testes and the gonads express female levels of DMRT1. If estrogen signalling is blocked in males heterozygous for functional DMRT1, the gonads develop as testes and express male levels of DMRT1. Similarly, when estrogen signalling is blocked in wild-type female embryos, DMRT1 levels increase and the gonads develop as testes.

It is well established that the epithelium of the left male gonad expresses ER and has the potential to form an ovarian cortex if exposed to estrogen (26). In the mixed-sex chimeric gonads analysed here, ER was clearly activated in both male and female epithelial cells. However, in the case of the somatic medullary cells, ER was only activated in the female medulla. It is noteworthy that, in the mixed-sex chimera, the female medulla and both the female and male epithelium express low levels of DMRT1, while the male medulla expresses high levels of DMRT1. It is also interesting that while epithelial cells of the male gonad show a paracrine response to estrogen, cells of the male medulla do not respond. Thus the differentiation of gonad epithelial cells is hormone dependent, whereas the differentiation of medullary somatic cells is cell autonomous, which can explain why male and female somatic cells in the medulla can retain their own sex identity (CASI) in the mixed-sex chimeric gonads (Figure 2) (17).

It is clear that higher levels of DMRT1 transcription in male gonads are closely linked to testis development. It is generally assumed that this sex difference is a result of a lack of dosage compensation of this Z-linked gene. In addition, a recent gene-editing study suggests that two functional copies of DMRT1 are required to inhibit FOXL2 expression and estrogen synthesis in the male. However, it is also clear, that if estrogen signalling is blocked, a single functional copy of DMRT1 is sufficient to promote testis development. The current study suggest that, although transcription levels are higher in males than in female, in the initial stages of gonadal development, DMRT1 protein levels are equivalent in male and female gonads – presumably as a result of translational or post-translational control. In addition, our findings suggest that the sex difference in DMRT1 protein levels seen in later stages of gonadogenesis, is due to a reduction in DMRT1 levels in female gonads – and that this is due to estrogen acting in a cell autonomous (autocrine) manner (Figure 7).

**Figure 7.**
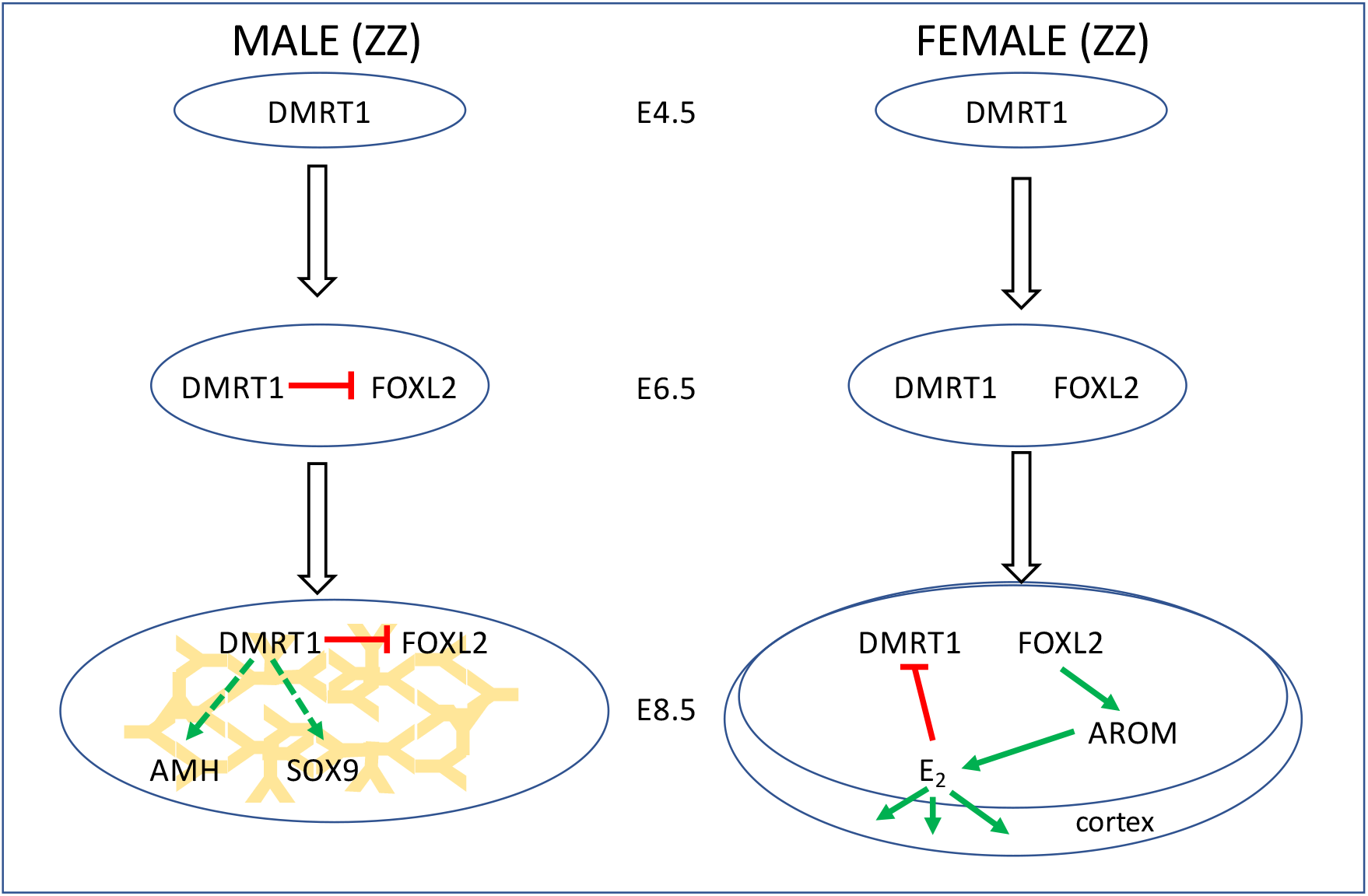
Schematic illustrating gene expression and molecular interactions in the developing testis and ovary. During the bipotential phase of gonadal development, the somatic cells of the medulla in both male and female embryos, express high levels of DMRT1 protein. During the differentiation phase, medullary cells of the male gonad continue to express high level of DMRT1 protein, inhibiting FOXL2 expression, and promoting sex cord formation and inducing AMH and SOX9 expression. In the female medulla, FOXL2 expression induces aromatase expression which enables the conversion of androgen to estrogen. Estrogen promotes cortex formation and suppresses DMRT1 expression.

Our data suggests that DMRT1 initiates gonadal development and induces medullary growth and primitive sex cord formation in both male and female chick embryos. Our findings also support the theory that the developmental outcome (testes or ovary formation) is a result of different levels of DMRT1 protein in the developing gonads of male and female embryos. However, it appears that the sex difference in DMRT1 protein levels is due to a reduction in DMRT1 protein expression in female gonads rather than simply a result of elevated levels of transcription in male embryos. In male gonads, DMRT1 promotes Sertoli cell differentiation and sex cord formation, while in the female gonad, estrogen signalling both stimulates cortex formation and suppresses DMRT1 expression, inhibiting sex cord formation in the medulla.

Further research is required to establish why DMRT1 does not inhibit FOXL2 expression in female gonads.

## Acknowledgements

We thank the National Avian Research Facility at the Roslin Institute for animal husbandry services and the provision of eggs and the BioImaging facility for technical assistance. We thank M. A. Hattori for kindly providing VASA antibody, and S. Guioli, R. Lovell-Badge and C Smith for DMRT1 antibodies.

## Authors Contributions

MC devised approach and obtained funding. DZ, LL, SN, JI and XY performed manipulations and analyses. MC, DG and MJM supervised procedures. MC and DZ wrote manuscript and all authors contributed to editing and revision.

## Materials & Methods

### Tissue collection

Freshly laid fertile eggs were incubated blunt side up, at 37.5°C, in 60 % humidity, with rocking (one rotation per 30 minutes) for the desired incubation period.

Eggs were removed from the incubator at the required developmental stage and embryos were carefully removed, sacrificed according to Home Office Schedule I procedures and the gonads dissected and processed for further analysis. Gross morphology of gonads was recorded using a Zeiss Axiozoom Microscope (Carl Zeiss AG).

For RNA analysis, gonads were dissected, placed in PBS, and any remaining mesonephric tissue removed. Gonads were snap-frozen in 10 μL of RNA-Bee (AMS Biotechnology) until RNA extraction. For Western analyses, gonads were collected into 100 μL of RIPA buffer (Thermo Fisher Scientific). For immunostaining, gonads+mesonephroi were placed in 4 % paraformaldehyde. A small portion of embryonic wing tissue was collected and used to determine genetic sex (36).

### Fadrozole Treatment

E2.5 embryos were injected with the aromatase inhibitor Fadrozole (FAD) (21–23) A small hole was made in the blunt end of the egg and 0.1 mg FAD dissolved in PBS was injected into the air sac. The eggs were then sealed and reincubated until gonads samples were collected as described.

### Protein Extraction and Western Blotting

Total protein was extracted from gonads into RIPA buffer (Thermo Scientific). For E4.5/E5.5/E8.5 embryos, ten/five/three pairs of gonads from the same gender were pooled, respectively (5 pools generated for each sex and for each stage).

Relative levels of DMRT1 protein in individual samples were estimated using an ‘Odyssey-based Western Blot Analysis’ method (http://biosupport.licor.com). Briefly, protein samples were subjected to electrophoresis on Bis-tris gels and then transferred to PVDF membranes. Membranes were blocked in Intercept Blocking Buffer for 1 hour (LI-COR Biosciences) and incubated overnight with primary antibodies; rabbit anti-DMRT1, rabbit anti-γ-tubulin, T3559, Sigma. Following incubation with fluorescently-labelled secondary antibodies, the membranes were imaged using an Odyssey imaging system. Scanned images were analysed with ‘Image Studio Lite Ver 5.2’ software.

### Quantitative Real Time PCR

Individual gonad pairs were homogenized in RNA-bee (AMS Biotechnology) and the lysate was loaded onto a Direct-zol RNA Microprep RNA extraction column (Zymo Research) and DNase-treated as per the manufacturer’s protocol. First-strand cDNA was synthesized using the ‘First-strand cDNA synthesis kit’ (GE Healthcare) according to the manufacturer’s instructions. PCR reactions were optimised to meet efficiencies of between 95 % and 105 % across at least a 100-fold dilution series. QPCR reactions were performed using a Stratagene MX3000P qPCR system (Agilent Technologies). Relative RNA expression levels were calculated using the 2^-ΔΔCt^ method} (37).

### Immunohistochemistry

Immunohistochemistry was carried out according to the protocol described by Stern^43^. Gonads were fixed in 4 % paraformaldehyde for 2 hours at 4°C. Tissues were equilibrated in 15 % sucrose/0.012M phosphate buffer overnight, embedded in 15 % sucrose plus 7.5 % gelatin/0.012M phosphate buffer (pH 7.2) and snap frozen using isopentane. Ten micrometer (10 μm) thick sections were cut on a cryostat (OTF 5000 Bright Instruments) and mounted on Superfrost Plus slides (Thermo Fisher Scientific). Slides were de-gelatinised for 30 min in PBS at 37°C and blocked in PBS containing 10 % donkey serum, 1 % BSA and 0.3 % Triton X-100 for 2 hours at room temperature. Incubation with primary antibodies was carried out overnight at 4°C, followed by washing four times in PBS containing 0.3 % Triton X-100, and incubation with secondary antibodies for 2 hours at room temperature. After washing four times in PBS containing 0.3 % Triton X-100, the sections were treated with Hoechst nuclear stain solution (10 μg/ml) for 5 min. Imaging was carried out using a Leica DMLB Upright Fluorescent microscope (Leica Camera AG).

### Generation of chimeric gonads

Embryo transplantation was carried out as described previously (17).

