## Supplementary Figure 1 for "Estrogen suppresses DMRT1 expression during ovarian development in the chicken"

**Suppl.Fig. 1. Chicken DMRT1 and transcript isoforms**

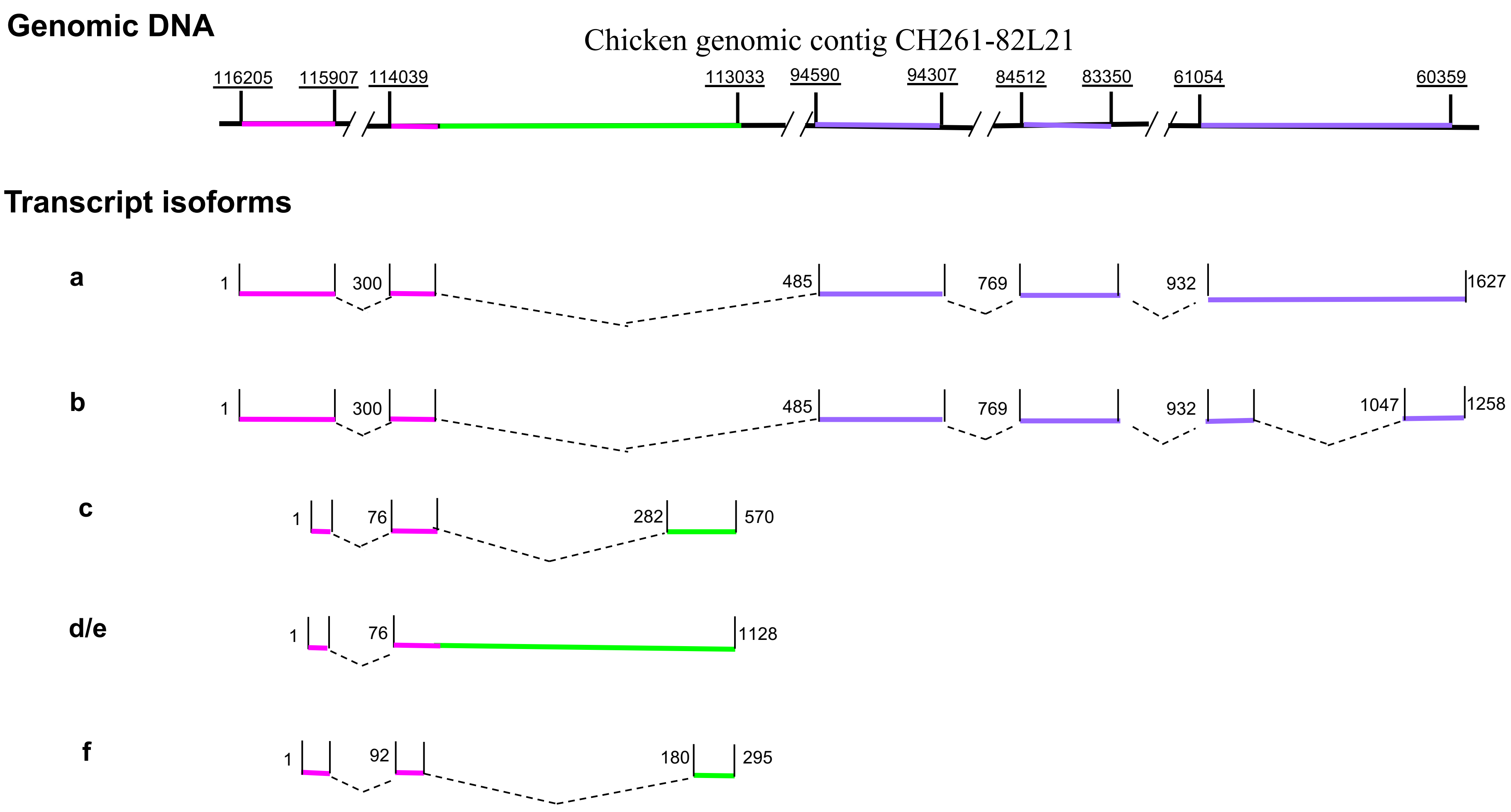


Relationship between genomic sequence and novel DMRT1 transcript isoforms.
DMRT1 locus spans over 65 kb on the chicken Z chromosome.
